# Age-Related Changes in the Neural Dynamics of Speech Production Following Noninvasive Brain Stimulation

**DOI:** 10.64898/2026.07.20.739631

**Authors:** Fatemeh Tabari, Joel Isaac Berger, Karim Johari

**Author notes:** Corresponding to: Karim Johari, Ph.D., Assistant Professor, Department of communication sciences and disorders, Louisiana State University, Baton Rouge, LA.

## Abstract

**Introduction:** Non-invasive brain stimulation is a promising technique to restore neural function in older adults by enhancing cortical excitability. In the present study, we examined the behavioral and neural effects of three stimulation protocols over the supplementary motor area on speech production in older adults. These protocols included fixed-frequency and personalized high-definition transcranial alternating current stimulation (HD-tACS) and high-definition transcranial random noise stimulation (HD-tRNS). We also compared the effects of personalized HD-tACS and tRNS between younger and older adults. Two groups, older adults (mean age: 69 y ± 8.18) and younger adults (mean age: 19.87 y ± 1.54), participated in this study. Older adults attended a four-session experiment including sham, fixed-frequency beta-HD-tACS (17 Hz), personalized beta-HD-tACS, and HD-tRNS. Younger participants also followed a similar study design but without fixed-frequency beta-HD-tACS. Following 25 minutes of stimulation, participants were asked to complete a speech production task while their EEG activity was recorded. Both personalized tACS and tRNS significantly reduced reaction times compared to sham in younger and older adults. In older adults, personalized tACS and tRNS both effectively modulated speech motor network but exhibited distinct spatial profile. While personalized tACS produced more focal modulation, tRNS elicited broader modulation across distributed neural networks and engaged multiple oscillatory frequencies. Broadband and personalized stimulation approaches (personalized tACS and tRNS) were more effective than fixed-frequency beta-tACS in modulating behavioral and neural responses measured throughout a distributed speech-motor network in older adults.

## 1. Introduction

Normal brain aging is characterized by a variety of changes including, but not limited to, cellular, molecular, metabolic, neural, and structural alterations such as loss of grey matter and white matter volume [1]. Neural changes are accompanied by reduced motor cortex plasticity and cognitive reserve [2, 3]. Such age-related declines affect daily life functioning by deteriorating action initiation, motor control and coordination, and even learning new motor skills [4].

Notably, older adults show more variable reaction times during motor tasks, attributed to slower neural processing, reduced motor function (preparation and execution), impaired cognitive control, sensory deficits, and recruitment of additional neural resources [5, 6]. The overactivation of additional prefrontal and motor regions is considered a compensatory mechanism to overcome age-related structural changes and maintain motor execution through reorganization and redistribution of functional networks [4]. Mattay et al. (2002) reported more pronounced bilateral and contralateral activation in the sensorimotor regions and cerebellum, the lateral premotor area, and the supplementary motor area (SMA) in aging brains, even for a simple button-press motor task [7].

Moreover, converging evidence has reported alterations in neural oscillatory patterns in aging brains, such as altered beta-band (15-30 Hz) oscillations during movement preparation and execution or modulation of the alpha (9-13 Hz)-theta (4-8 Hz) bands in cognitive-motor integration, resulting from neurotransmitter system decline, structural changes, and compensatory reorganization [8-11]. Park et al. (2025) found a progressive decrease in alpha-band power and increase in beta-band power extending from the parietal to the sensorimotor region during the resting state in older adults compared to their younger counterparts [12]. Theta band power decreases in frontal regions were also reported as a neurophysiological signature of aging cognitive decline [13]. The findings of a multicenter EEG study showed a decrease in resting-state magnitude of the posterior delta-band (1-3 Hz), which linearly correlated with age [13]. However, such alterations in oscillatory mechanisms are task- and region-dependent. For instance, the results of a MEG study in older adults showed increases in beta-band power during rest and decreases during a hand movement task, specifically in the contralateral primary motor cortex (M1) [14]. A strong decrease in beta-band power in frontal regions during a force modulation task [15] and decreases in the left posterior temporal and inferior parietal cortex during spoken word production were also reported [16].

An emerging body of research suggests that non-invasive brain stimulation, such as transcranial electrical stimulation (tES), can improve motor and cognitive functioning by enhancing neural plasticity and cortical excitation [17, 18]. tES encompasses multiple stimulation techniques, including transcranial direct current stimulation (tDCS), alternating current stimulation (tACS), and random noise stimulation (tRNS) [19]. Several studies have robustly supported the beneficial effects of anodal tDCS over various cortical regions, such as M1, dorsolateral prefrontal cortex, and the inferior frontal gyrus, across various tasks, including motor tasks (such as finger tapping), discourse production, verbal fluency, and naming in older adults [20-22]. Anodal tDCS over the SMA also improved motor coordination and postural stability in older participants [23]. Despite the promising potential of tDCS, a high degree of inter-individual variability in response to stimulation has raised some concerns regarding the replication of results for older individuals [24]. Notably, age-related anatomical changes may attenuate the strength of the tDCS-induced electric field [25], highlighting the need for frequency-specific and individualized stimulation in older adults.

tACS is another non-invasive stimulation technique that modulates endogenous brain oscillations, potentially through entrainment induced by exogenous alternating currents [26]. Entrainment can be applied at a fixed (e.g., 20 Hz) or an individualized frequency [27]. Personalized tACS has shown potential in improving cognitive and motor function by aligning the stimulation with an individual’s neural profile in healthy adults and normalizing disrupted neural oscillations in those with neurological disorders, particularly Parkinson’s disease [28, 29]. tRNS is also a variation of tACS which delivers weak alternating electrical currents at random and constantly changing frequencies, typically spanning from 100 to 640 Hz [30]. tRNS has shown improvements in sensorimotor and cognitive functioning in younger adults [31, 32]; whereas its effects on older adults remain inconsistent, with evidence suggesting that these effects may depend on age and task-specific factors [33].

While the facilitatory effects of personalized high-definition tACS (HD-tACS) have been demonstrated in young adults [34], its applicability to the older adults remains largely unexplored. Given the age-related alterations in anatomical profiles and oscillatory dynamics, findings from younger cohorts may not be generalizable to older adults. Hence, it is critical to examine whether personalized stimulation can optimize neural functioning in older populations. Our previous study showed that personalized beta-band, SMA-targeted stimulation could effectively modulate neural oscillations in the delta-band (1-3 Hz) and improve behavioral performance (i.e., faster reaction times) in young adults [34]. The current study investigated the effect of fixed-frequency and personalized HD-tACS and HD-tRNS over the left SMA – a neural hub for initiation, coordination, execution, and control of speech motor functions [35] – on speech production in healthy older adults (age > 60 years) and compared results with a younger cohort. We hypothesized that (a) SMA-targeted HD-tACS tuned to each individual’s dominant beta frequency would facilitate speech motor control and underlying neural oscillations, and (b) older adults would derive greater benefit from personalized stimulation than younger adults, given their lower baseline neural efficiency and age-related oscillatory alterations. The beta-band (15-30 Hz) was selected for personalization purposes because it is a dominant neural signature in motor preparation, execution, and motor control [36, 37]. This study included new data from an older group of participants, along with data from a younger cohort that were previously published [28, 34]. The integration of both groups enables a direct comparison of age-related differences in response to personalized and non-personalized stimulation protocols at the neural and behavioral levels.

## 2. Methods

The study comprised twenty older adults (mean age: 69 y ± 8.18; age range: 60–84; 15 women) and twenty-two younger adults (mean age: 19.88 y ± 1.54; age range: 18–27; 17 women). All participants were informed about the purpose of the study and written informed consent were obtained. Older participants were financially compensated, and younger participants were compensated with either course credit or monetary payment for their time. This study was approved by the Institutional Review Board of Louisiana State University (IRB approval number: IRBAM-21-1019).

Inclusion criteria included being a native English speaker, right-handed, having normal or corrected-to-normal vision and hearing, and no history of neurological or communication disorders. The minimum age was 60 years for older adults and 18 years for younger adults. For older individuals, cognitive status was assessed using the Mini-Mental State Examination (MMSE) and Frontal Assessment Battery (FAB), with all participants scoring within the normal range for healthy older adults (MMSE = 29.5 ± 0.57; FAB = 17.4 ± 0.56).

### 2.1. Stimulation Configuration

The configuration of electrodes was determined using HD-Target and HD-Explore software (Soterix Medical, New York, NY). The MNI152 template was used to optimize the intensity and focality of current flow to the left SMA. MNI coordinates and corresponding electric field intensities are presented in Table 1. Figure 1A illustrates the offline modeling of the current flow.

**Figure 1.**
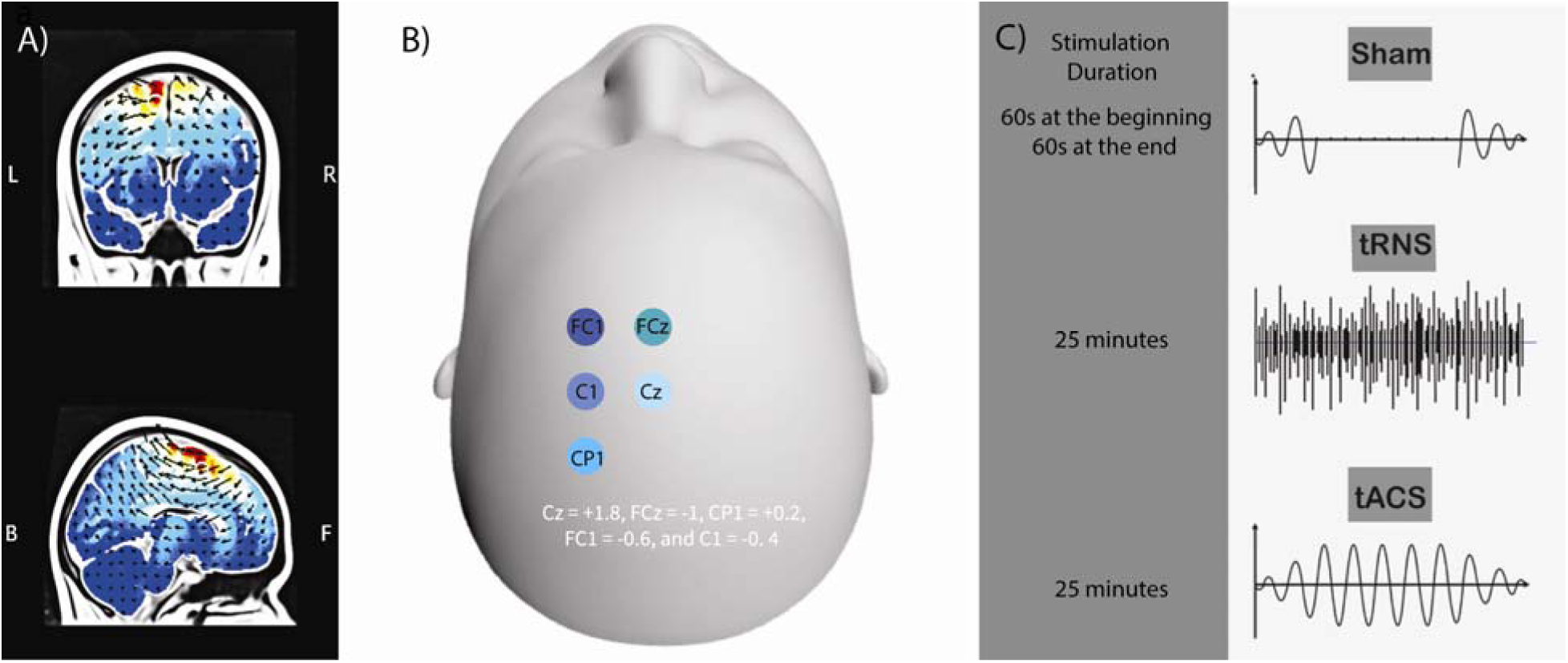
(A) The modelled current flow of HD-tACS obtained using the HD-Target and HD-Explore software (Soterix Medical, New York, NY) (B). Placement of 4 × 1 HD-tACS electrodes according to the 5-10 standard EEG montage with values of Cz = +1.8, FCz = -1, CP1 = +0.2, FC1 = -0.6, and C1 = - 0.4. (C) Demonstration of tES protocols and the stimulation duration.

**Table 1.** Field intensity (v/m) within the left SMA obtained by offline modeling based on configurations in Fig. 2 A and 2B.

| MNI Coordinates |  |  | Filed Intensity<br>(v/m) |
| --- | --- | --- | --- |
| x | y | z |  |
| -11 | 1 | 56 | 0.16 |
| -11 | -4 | 60 | 0.15 |
| -12 | -1 | 57 | 0.17 |
| -8 | -6 | 67 | 0.15 |
| -8 | -11 | 64 | 0.16 |

HD electrodes were positioned on the scalp using the 5-10 international montage. Conductive gel was applied between the skin surface and the electrodes to lower the skin-electrode impedance (<15 kΩ). HD-tACS, HD-tRNS, and sham stimulations were administered using the M×N-9 HD-tES stimulator (Soterix Medical Inc., NY, USA).

The participants were blinded to the stimulation condition. They were instructed to report any pain and unpleasantness 30 seconds after stimulation onset, at the midpoint, and 30 seconds before the end of the stimulation. All participants tolerated the stimulation well, and no adverse effects were reported.

### 2.2. Experimental Design

The participants completed testing in a sound-attenuated booth. Visual stimuli were presented on a monitor at a viewing distance of 70 cm. Each stimulation session (including sham, in which 23 minutes of the session involved no stimulation, without explicit instruction to the subject that this was the case; see section 2.4) lasted 25 minutes. EEG and stimulation were not performed concurrently to avoid stimulation-induced artifacts. Following stimulation and a 20-minute preparation period, EEG data were recorded for 25 minutes. The participants performed an interleaved speech production (vowel vocalization) and limb movement (button press) with concurrent recording of behavioral and neural data. Each trial started with a 1000-1500 ms fixation period, followed by a blank screen for 500-1000 ms. Then a cue appeared for a duration of 1500-2000 ms, followed by a blank screen lasting 500-1000 ms. The speech cues required participants to produce and sustain a steady vocalization of one of the long vowels (/aa/, /ee/, or /oo/) or the short vowel (/o/) upon the appearance of the green circle and cease vocalization upon its disappearance. The limb cues required them to press the corresponding keyboard key ( ) with their right index finger in response to directional arrows (up, down, right, or left; 1500-2000 ms) and to maintain the press until the circle disappeared (Figure 2).

**Figure 2.**
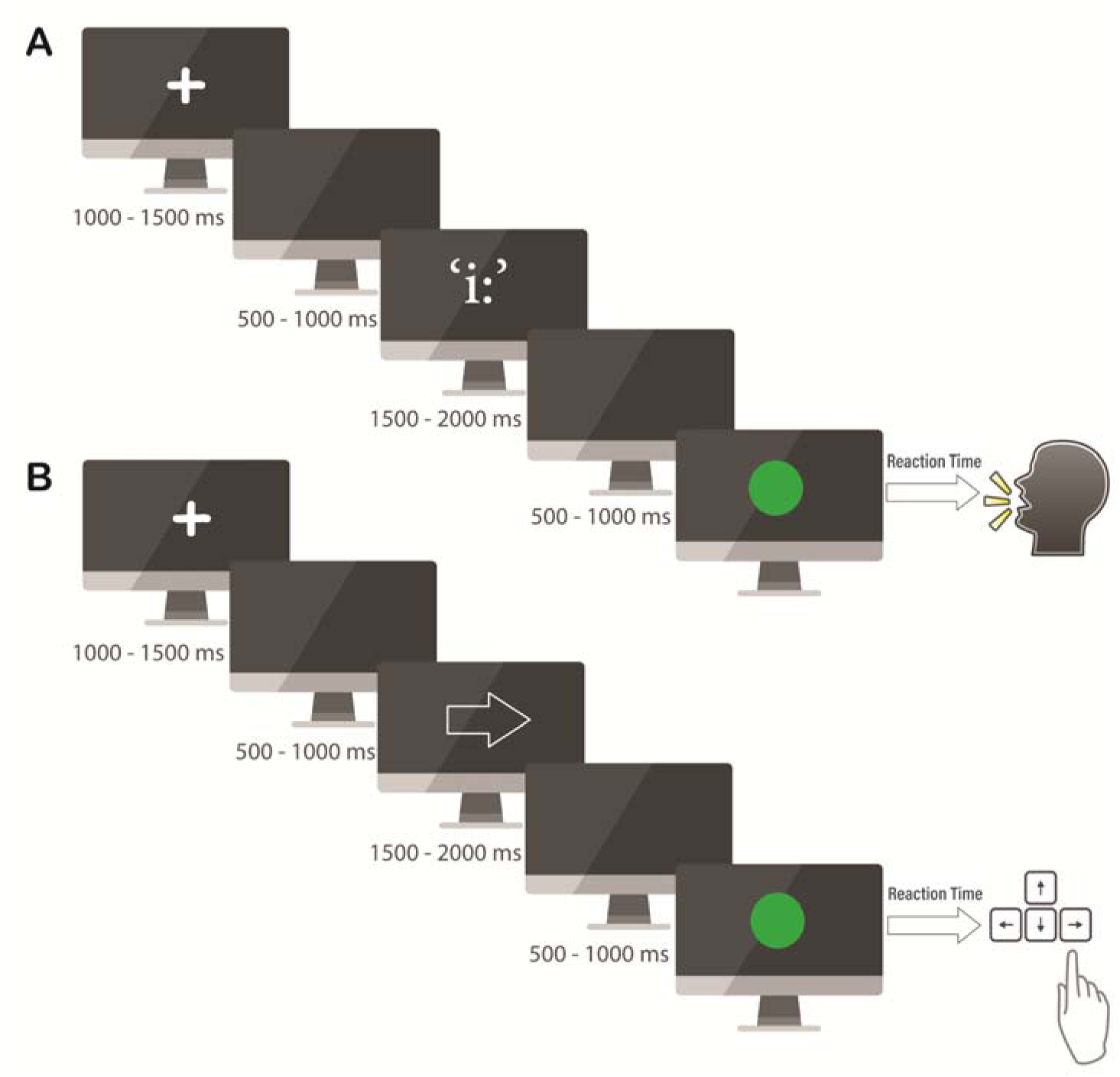
Visual representation of the study design schematic for Speech (A) and Finger Press (B) tasks.

The experimental task was implemented using a custom-made MATLAB R2022a (MathWorks Inc.) script in Psychtoolbox [38]. The script generated event markers recorded behavioral responses (reaction times and accuracy), and synchronized these with EEG data. A total of 200 trials were collected, with 100 trials for each modality (speech and limb movement, i.e., finger press). The limb cues were initially included in our experimental protocol to examine whether the stimulation effects on the SMA - as a shared network for motor functions - also extended across hand motor domains and were not specific to speech, consistent with our previous studies in young participants [28, 34]. In the present study, we specifically restricted our analysis to speech production to focus on stimulation-induced neural and behavioral changes within the speech motor network; therefore, the data collected from limb tasks fall outside the scope of this study.

Participants’ acoustic data were captured using a Focusrite digital audio recorder (Focusrite Scarlett 2i4, High Wycombe, UK) and a high-quality headset condenser microphone. A headset microphone was used to ensure a constant mouth-to-microphone distance of ∼5 cm and minimize head movement noise.

### 2.3. EEG Acquisition

EEG data were recorded using 64-channel BrainVision actiCAP active electrodes (Brain Products GmbH) following the 10-5 international montage at a sampling rate of 500 Hz. The ground electrode and online reference were positioned at FPz and Fz, respectively. To ensure optimal signal quality, conductive gel was applied to lower electrode-skin impedance (< 5KΩ). EEGLAB software [39] was used to preprocess the data. EEG signals were downsampled to 200 Hz, high-pass filtered at 1 Hz to remove low-frequency components (e.g., DC shifts), and visually inspected for noisy electrodes, which were then interpolated. Independent component analysis (ICA) was subsequently applied to detect and remove physiological or external artifacts (e.g., eye blinks, muscle activity, and line noise) [40].

### 2.4. Stimulation Design

In a within-subject study design, participants completed four stimulation conditions, including three active stimulation sessions and one sham condition. For older adults, active stimulation included Tuned-to-Speech [TtS]-HD-tACS, Tuned-to-Young [TtY]-HD-tACS, and HD-tRNS stimulation (hereafter referred to as TtL, TtY and tRNS, respectively) with a 25 min, 2 mA stimulation applied over the left SMA. The order of conditions was counterbalanced across the participants; however, TtS and tRNS were never administered as first sessions, as both required the extraction of peak frequency from the sham condition. The same electrode montage and protocol was applied for sham condition. No continuous stimulation was applied during the sham session; instead, a brief 60-second ramp-up and ramp-down stimulation was delivered at the beginning and the end of the 25-minute period to induce sensations similar to active stimulation. To specify the frequency of the TtS condition, EEG data collected following the sham session were used to extract each individual’s peak frequency of beta-band within the left SMA for speech production. A similar design was used for younger adults, comprising three active stimulation sessions : TtS, Tuned-to-Limb [TtL; tuned to the beta peak frequency extracted from button press tasks], HD-tACS, HD-tRNS stimulation– and a sham session (See Tabari et al., 2026 for details). In the present study, only TtS, tRNS, and sham conditions of the younger group were included, because these were common in both older and younger groups. The TtL condition was not added to the analysis because the primary goal of the study was to investigate speech-motor responses to stimulation rather than limb motor performance.

### 2.5. Peak Frequency Extraction

After preprocessing, sensor-space EEG data obtained during the sham session were projected into source space to identify each individual’s peak frequency at the cortical vertex closest to the stimulation target; based on Euclidean distance to the MNI coordinates provided by the modeling software. The inverse operator was estimated using the MNE-Python toolbox [41] based on the dSPM method. Data were projected onto an average template brain (fsaverage) and filtered between 0.1 and 50 Hz. Source-localized EEG data were subsequently epoched for the speech task within a time window from -1000 ms to 0 ms prior to vocalization and the average source time course was extracted. Finally, a fast Fourier transform was applied to these data in MATLAB to estimate the maximal frequency within the beta range (13-30 Hz) for each individual. Participants’ stimulation frequencies for TtS were defined based on the obtained values.

The mean peak frequency of beta band for TtS in older adults was 16.5 Hz (range = 13-24 Hz) and 17 Hz (range=14-28) for younger adults. The TtY frequency was set at 17 Hz, corresponding to the mean beta peak frequency of the younger cohort in our prior study [28]. The frequency of the sham condition was fixed at 15.5 Hz, informed by previous findings showing the differential modulation in low beta (13-18 Hz) and high beta (18-25 Hz) for a similar task [42]. Accordingly, 15.5 Hz was chosen as it represents the midpoint between 13 and 18 Hz. The frequency of tRNS was set as 30 Hz, but for individuals with peak frequencies closer to 30 Hz, HD-tRNS was set as 35 Hz to maintain at least a 10-Hz difference from the TtS condition, allowing for an unpredictable amplitude while inducing similar scalp sensation to the tuned stimulation.

### 2.7. Neural Data Analysis

EEG data were preprocessed, analyzed, and visualized using EEGLAB software [43] and MATLAB. For frequency analysis, data were segmented into epochs of -1500 ms to 1000 ms after the onset of speech. The spectral power of neural data in the range of 1-30 Hz (1 Hz resolution) was computed using Morlet wavelets. Neural oscillatory power time-locked to the onset of vocalization was computed for each trial as the squared magnitude (i.e., absolute value magnitude) of the complex wavelet coefficients. The following formula was used to calculate the power changes in EEG activity for each frequency on a per-trial basis:

Power[dB]=10×log10_(Response Period/Baseline Period)_

with the baseline set at 1000 to 0 ms before the onset of the vocalization to minimize contamination from anticipatory speech-motor preparation. Data were log-transformed to approximate normality.

Our statistical analysis was limited to the beta-band (15-30 Hz) and the alpha-band (9-13 Hz), as the beta-band corresponds to the stimulation frequency and the alpha-band was selected because of its sensitivity to age-related changes [43]. The choice of frontocentral and central electrodes was informed by previous studies following similar tasks [28, 44].

### 2.6. Statistical Methods

Speech reaction times (RT) were calculated as the temporal difference between the visual stimuli and the point at which the participant’s voice intensity surpassed 10% of its peak amplitude. The measures were submitted to a linear mixed-effects model with age group (Young, Aged) and stimulation (Sham, TtS, TtY, and tRNS) as fixed effects and subject as a random intercept. The differences between the groups and stimulation conditions were examined using post hoc comparisons of estimated marginal means (EMMs) with Bonferroni correction. Behavioral data were analyzed using R software (R Core Team, 2013).

For neural analysis, the spectral power values associated with the beta (15-30 Hz) and the alpha (9-13 Hz) bands were extracted for the time window of -500 to 500 ms. The values were subsequently submitted to cluster-based permutation testing to detect significant temporal differences across stimulation conditions, within and between groups [45]. Cluster-level significance was set at α = 0.05 using 5000 permutations. This method offers an unbiased and strong approach for evaluating the spectral power.

## 3. Results

### 3.1. Behavioral Results

The results of the study are presented at two levels of analysis: within-group and between-groups comparisons. Descriptive statistics for speech production reaction times (RTs) are presented in Table 1 across three stimulation conditions and two age groups. Sham, Tuned-to-Speech (TtS), and tRNS are stimulation conditions common to both younger and older adult groups. Tuned-to-Young (i.e., the condition in which the stimulation was set to the mean peak frequency of young adults) is reported only for older adults. TtY was included as a comparison condition to determine whether the stimulation-induced effects were driven by individualized, age-specific oscillatory characteristics or could be achieved using a fixed beta frequency derived from younger adults.

#### 3.1.1. Between-Group Comparisons of Younger and Older Adults

Table 2 shows descriptive statistics of speech production reaction times (RTs) of each stimulation condition for younger and older adults. Although the standard deviations (SD) were numerically larger in the older group, Levene’s tests were conducted to examine the homogeneity of variance. The tests revealed no significant differences across all conditions (all *p* > 0.23), indicating that the variance was comparable between age groups.

**Table 2.** Group means (milliseconds) and standard deviations (SD) of speech RTs in young and older adults across stimulation conditions.

| Age Group | Stimulation | Mean (ms) | SD |
| --- | --- | --- | --- |
| Young Adults | Sham | 430.848 | 118.957 |
|  | TtS | 361.242 | 85.86 |
|  | tRNS | 373.121 | 85.961 |
|  | TtY | 490.221 | 146.159 |
| Older Adults | Sham | 527.597 | 140.061 |
|  | TtS | 454.572 | 129.521 |
|  | tRNS | 438.772 | 106.158 |

The results of a linear mixed-effects analysis revealed a significant main effect of age group (*F*(1, 44.13) = 7.87, *p* = 0.007), indicating that older participants exhibited slower responses overall than younger participants. There was also a significant main effect of stimulation, *F*(2, 86.32) = 26.35, *p* < 0.001, indicating that RTs differed across stimulation conditions. The age group × stimulation interaction was not significant, *F*(2, 86.32) = 1.05, *p* = 0.355, suggesting that stimulation effects were similar for both age groups. Bonferroni-corrected post-hoc comparisons showed that younger participants were significantly faster than older participants in both sham (Mean Difference (MD) = 96.75 ms, *p* = 0.006, *d* = 0.75) and TtS conditions (MD = 92.00 ms, *p* = 0.008, *d* = 0.86). The difference in the tRNS condition showed a trend toward significance but did not reach the corrected threshold (MD = 65.76 ms, *p* = 0.056, *d* = 0.69). Figure 3A shows the comparison of speech RTs in young and older adults across the three common conditions.

**Figure 3.**
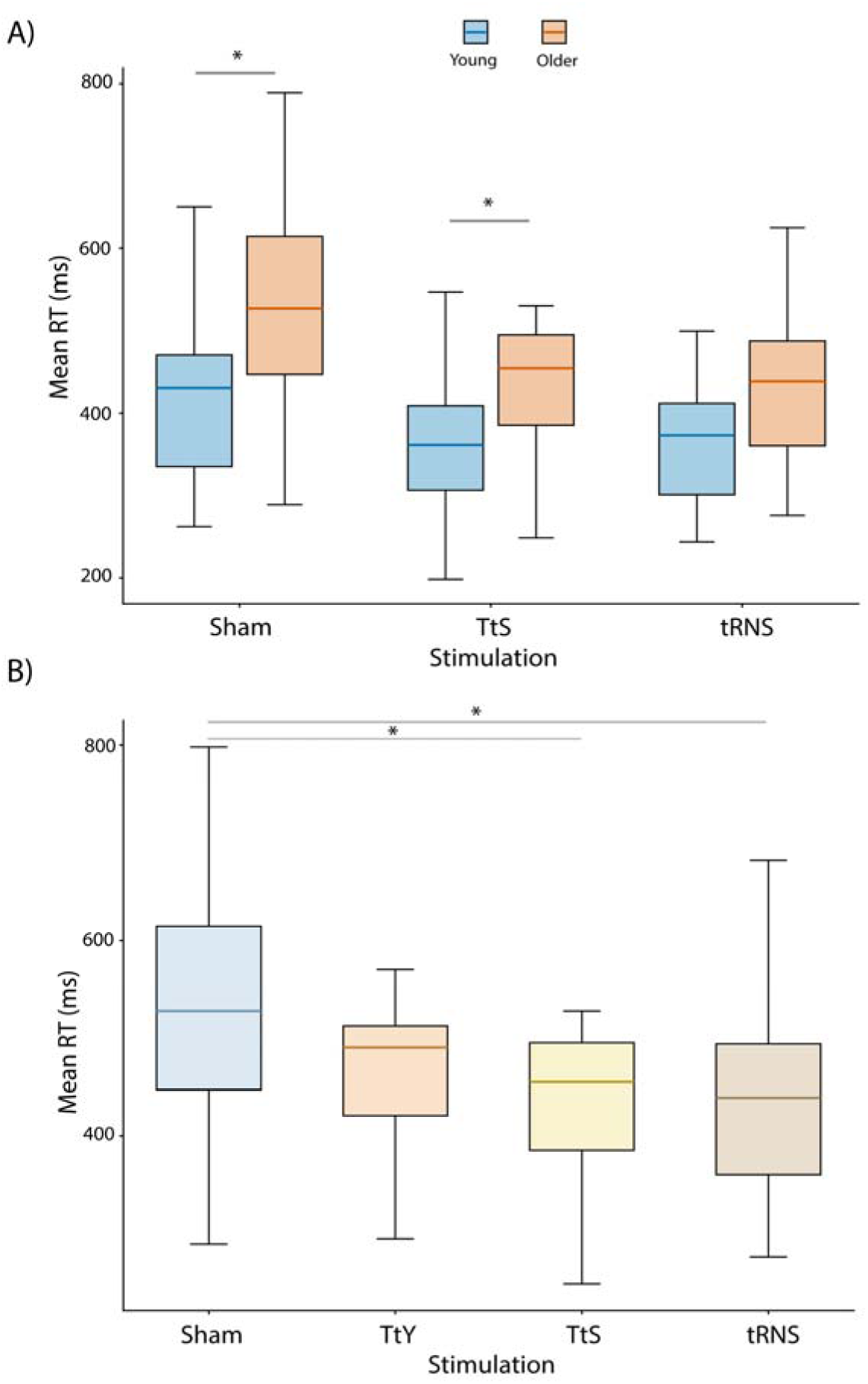
RTs in speech tasks (A) across three stimulation conditions (sham, Tuned-to-Speech [TtS], and tRNS) in younger and older adults, (B) across four stimulation conditions (sham, Tuned-to-Young [TtY], Tuned-to-Speech [TtS], and tRNS) in older adults. Box plots represent the interquartile range, horizontal colored lines indicate the means. Asterisks denote significant differences between groups (p < .05, Bonferroni-corrected).

#### 3.1.2. Within-Group Effects in Older Adults

A linear mixed-effects model with stimulation as a fixed-effect and participant as a random intercept revealed a significant main effect of stimulation on RTs in older adults, *F*(3, 60) = 7.95, *p* = 0.001. Bonferroni-corrected pairwise comparisons indicated that RTs were significantly faster in both the TtS (MD = 73.02 ms, *p* = 0.004) and the tRNS (MD = 88.82 ms, *p* < 0.001) conditions relative to sham. Unlike TtS and tRNS, TtY stimulation did not significantly improve RTs relative to sham after Bonferroni correction (MD = 37.37 ms, *p* = 0.418). No significant differences were observed between TtS and tRNS (*p* = 1.00) in speech RTs. Figure 3B shows the comparison of speech RTs across the four stimulation conditions in older adults.

### 3.2. Neural Results

#### 3.2.1. tRNS-Sham comparison

Among active stimulations, transcranial random noise (tRNS) elicited the broadest modulation in the alpha-band (9-13 Hz) and the beta-band (15-30 Hz) power during pre- and post-speech stimulus intervals in older adults, particularly over bilateral frontocentral, right frontocentral, and central electrode sites following tRNS stimulation. Such effects were not observed in younger adults. TtS elicited more localized prominent modulation in older adults, predominantly over right frontocentral electrodes.

##### Beta-band Power in tRNS-sham condition

Cluster-based permutation testing of the beta-band power revealed no significant group differences in the sham condition between young and older adults (*p* = 0.383) over the bilateral frontocentral electrodes (AFz, F1, F2, FCz, AF3, AF4). However, following the tRNS condition, significant younger-older adult differences were observed in a cluster spanning −555 to 120 ms (*p* = 0.003). In within-group comparisons, the older adult group exhibited a significant cluster from −355 to −125 ms (*p* < 0.001).

##### Alpha-band Power in tRNS-sham condition

In the sham condition, there was no significant difference between the young and older adult group in alpha-band power over the bilateral frontocentral electrodes (AFz, F1, F2, FCz, AF3, AF4; p = 0.370). In contrast, a significant group-difference cluster was observed from −625 to 20 ms (*p* = 0.012) in tRNS condition. Within-group comparisons showed no significant sham versus tRNS clusters in the young group (*p* > 0.5); whereas the aging group exhibited significant stimulation effects (tRNS versus sham), with a significant cluster spanning −630 to −355 ms prior to speech onset (*p* < 0.001).

In young adults, a significant difference was identified between sham and tRNS in theta band (4-8 Hz) power in a cluster extending from -880 to -310 ms (*p* < 0.001). Such a modulation effect was absent in older adults (*p* > 0.5).

Overall, older adults showed stronger power modulation around speech onset following tRNS stimulation, in the alpha (9-13 Hz) and beta-band (15-30 Hz) bands across the bilateral frontocentral electrodes compared to younger adults. Figure 4 depicts the time-frequency representations and log power profiles in both groups. Similar patterns were observed over the central and right frontocentral electrodes following tRNS stimulation supporting the primary findings. Specifically older adults exhibited stronger and broader alpha-beta modulation over central electrodes and stronger alpha-band modulation over the right frontocentral electrodes around the speech onset compared with younger adults (Supplementary Figures S1 and S2).

**Figure 4.**
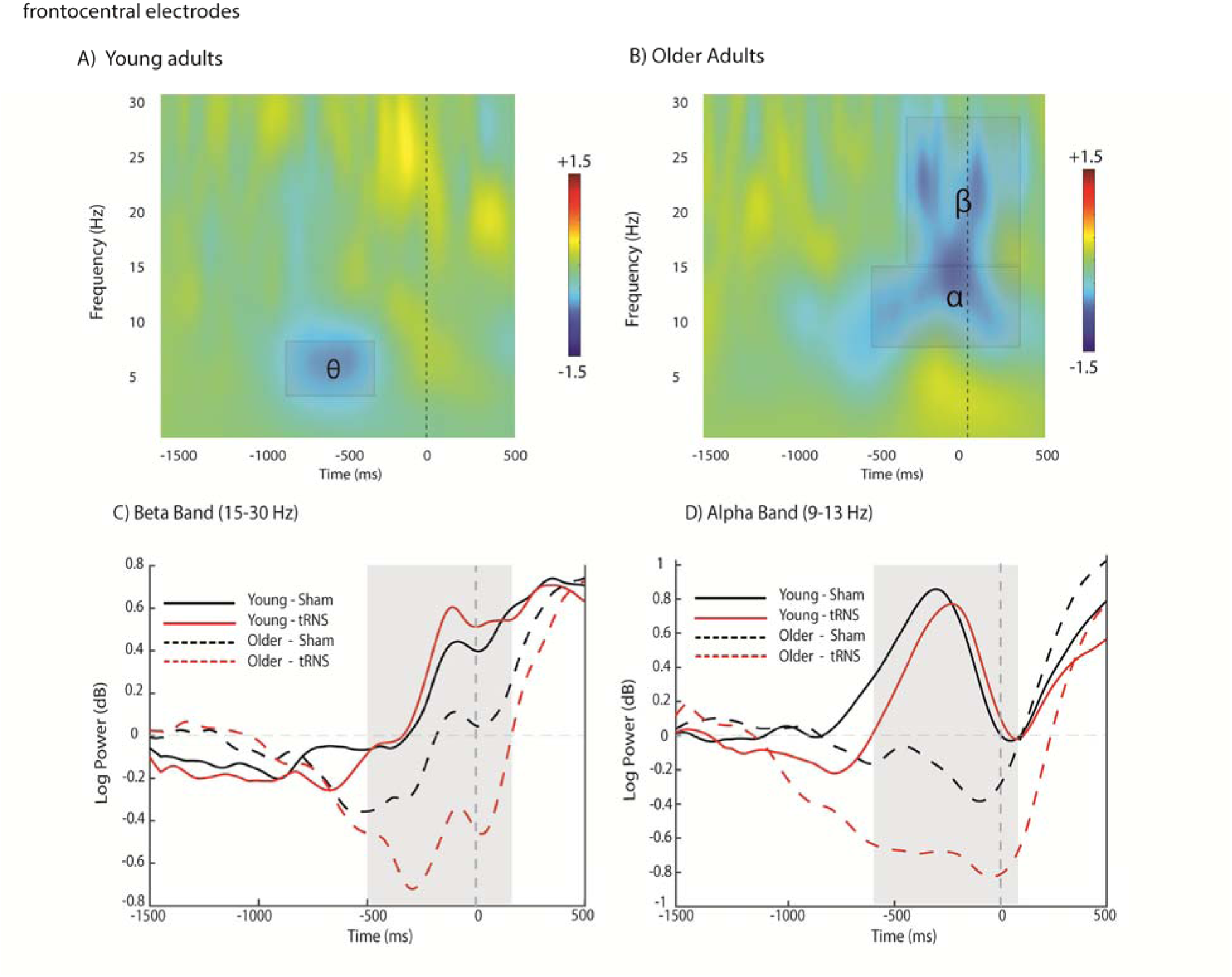
Neural activity following tRNS stimulation relative to sham averaged over the bilateral frontocentral electrodes (AFz, F1, F2, FCz, AF3, AF4) in young and older adults, across the time course of 1500 ms before to 500 ms after the onset of speech production. Time-frequency plots show power difference (tRNS-sham) for younger (A) and older adults (B). The black rectangles highlight the alpha/beta-band time-frequency cluster of interest in older adults and theta band (4-8 Hz) time-frequency cluster of interest in young adults, and the vertical dashed line at 0 marks the onset of vocalizing the speech sound. Overlaid temporal profiles of log power (in decibels: dB) show the beta-band (15-30 Hz: panel C) and alpha-band (9-13 Hz: panel D) time course for younger and older adults in tRNS (red) and sham (black) conditions. Solid lines represent the younger adults, and dashed lines represent the older adults. The shaded regions indicate the temporal clusters that reached significance in a cluster-based permutation analysis (between-group difference).

#### 3.2.2. TtS-Sham comparison

##### Beta-band Power (15-30 Hz) in TtS-sham condition

The analysis for the beta-band (15–30 Hz) power over the right frontocentral electrodes (AF8, F8, F4, F6, FC6) showed no significant group differences between young and older participants in the sham condition (*p* = 0.509); however, a significant younger-older group difference occurred from −650 to −130 ms (*p* = 0.012), with older adults exhibiting stronger beta-band modulation than younger adults following TtS stimulation. Within-group analyses did not reveal any significant differences in the temporal window of interest (*p* > 0.05).

##### Alpha-band Power (9-13 Hz) in TtS-sham condition

No significant younger-older differences were identified in the sham condition in the alpha-band (9–13 Hz) power within the right frontocentral electrode sites (AF8, F8, F4, F6, FC6). In contrast, a significant group-difference cluster was observed from −610 to 185 ms (*p* = 0.002) following the TtS condition.

Although there was an increase in alpha-band (9-13 Hz) modulation around the stimulus in young adults, a within-group comparison did not reach statistical significance (*p* > 0.05). However, the older adults exhibited a significant TtS-Sham cluster extending from −385 to 175 ms (*p* < 0.001).

In general, older adults exhibited stronger alpha modulation following TtS stimulation around the speech onset. Figure 5 shows time-frequency representations and temporal profiles of both young and older adult groups.

**Figure 5.**
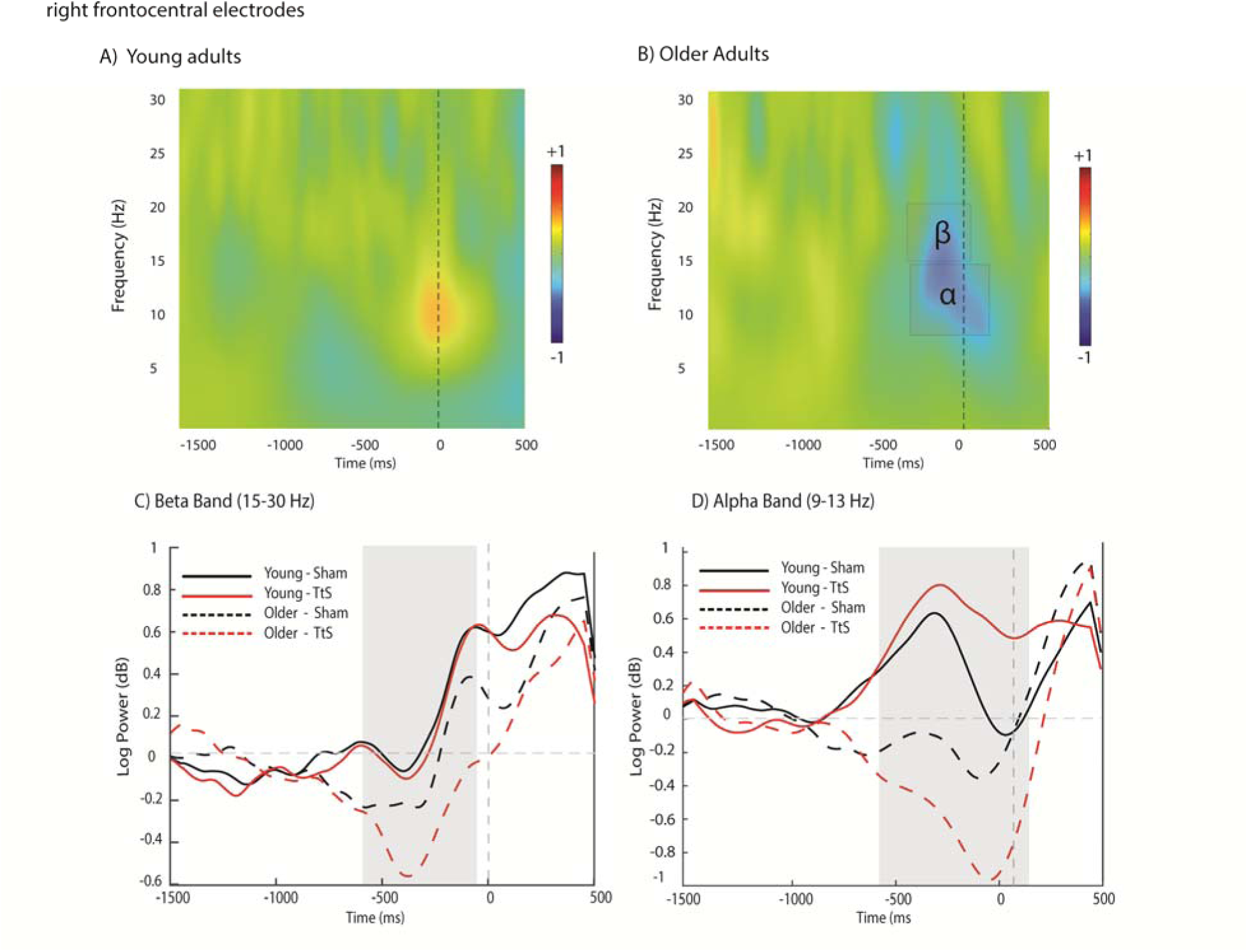
Neural activity following TtS stimulation relative to Sham over the right frontocentral electrodes (AF8, F8, F4, F6, FC6) in young and older adults, over the time course of 1500 ms before to 500 ms after the onset of speech production. Time-frequency plots show power difference (TtS-sham) for young (A) and older adults (B). The black rectangles highlight the alpha/beta-band time-frequency cluster of interest in older adults, and the vertical dashed line at 0 marks the onset of vocalizing the speech sound. Overlaid temporal profiles of log power (in decibels: dB) show the beta-band (panel C) and the alpha-band (panel D) time course for young and older adults in TtS (red) and sham (black) conditions. Solid lines represent young adults, and dashed lines represent older adults. The shaded regions indicate the temporal clusters that reached significance in a cluster-based permutation analysis (between-group difference).

Cluster-based permutation analysis identified no significant differences between young adults and older adults following TtS-sham stimulation in either central or bilateral prefrontal electrodes (*p* > 0.05) in the beta-band and alpha-band power around the speech onset.

#### 3.2.2. Within-group comparisons

Cluster-based permutation analysis of the beta-band (15–30 Hz) over bilateral frontocentral electrodes (AFz, F1, F2, FCz, AF3, AF4) in older adults revealed a significant difference between the sham and tRNS (−355 to −125 ms, *p* < 0.001), with no significant differences between sham and other active conditions (*p* > 0.05). tRNS also differed significantly from TtS (two clusters of −320 to −155 ms and −5 to 190 ms, *p* < 0.001) and TtY, yielding a brief significant cluster (spanning −290 to −275 ms, *p* < 0.001). In the alpha-band (9-13 Hz), however, all active conditions –tRNS (−630 to −355 ms), TtS (5 to 130 ms) and TtY (−545 to −395 ms)– differed significantly from sham (all *p* < 0.001), with no significant differences among the active conditions.

Additional within-group analyses were performed over central (Supplementary Figure S1) and right frontocentral (Supplementary Figure S2) electrode clusters to further characterize the spatial distribution of stimulation effects. Across the central electrodes (C1, C2, and Cz), there was no significant difference between any active stimulation condition and sham in the beta range. In contrast, in the alpha-band, both tRNS and TtS significantly differed from the sham (−180 to 100 ms and -30 to 6 ms, respectively: all *p* < 0.001). No significant differences were found among the active conditions in either band.

Over the right frontocentral electrodes (AF8, F8, F4, F6, FC6), the analysis showed significant differences in the alpha-band (9-13 Hz) for tRNS (−565 to −375 ms), TtS (−385 to 175 ms), and TtY (−425 to −390 ms) relative to sham. In the beta-band (15-30 Hz), a significant cluster was identified only for the tRNS compared to sham (−275 to −185 ms; all *p* < 0.001). No significant differences were observed among all other conditions in either frequency bands.

## 4. Discussion

This study is the first attempt to examine the modulatory effects of personalized beta HD-tACS and tRNS over the left SMA on speech-motor control in both younger and older adults. We investigated whether the task-specific personalized stimulation could improve speech production in older adults at both behavioral and neural level. This study expands on our previous research on younger adults group [28] by providing between-group (younger versus older adults) and within-group comparisons (older adults) to track age-related modulatory mechanisms of tACS/tRNS and their potential age-related benefits.

Between-group comparisons showed that older adults were significantly slower than younger adults at baseline and following TtS stimulation, with the difference reduced but approaching significance following tRNS. This finding is consistent with prior studies reporting age-related slowing of motor execution, likely due to decreased neural efficiency, impaired predictive timing mechanisms, and more extensive cortical recruitment during motor preparation and execution [46, 47].

At group-level (older adults), tRNS and TtS, but not TtY, could significantly reduce reaction times compared to the baseline (sham). These results align with previous works, supporting the modulatory effects of personalized tACS and tRNS [48, 49]. Wach et al. (2013) also suggested limited behavioral efficiency (slowing of motor functioning) following a fixed-frequency tACS (20 Hz), which is similar to our findings showing no behavioral improvement following TtY [50].

Consistent with the behavioral findings, TtY effects were also confined to early pre-motor planning (approximately 300 ms to 400 ms pre-onset interval), while both TtS and tRNS exhibited modulation extending from pre-motor to the execution phase, specifically in the alpha-band (approximately 500 ms pre- to 100 ms post-onset interval). This temporal pattern suggests that TtY (implemented as a fixed 17-Hz beta frequency) did not show evidence of task-relevant entrainment. The lack of neural and behavioral effects of fixed-frequency tACS reflects its inherent limitation in accounting for individuals’ variability in oscillatory patterns (especially in the older brain) and the multi-frequency nature of speech motor control.

Personalized beta stimulation (TtS) significantly improved speech performance and modulated speech-motor activity in older adults; however, its neural effects were more spatially localized, predominantly over right frontocentral electrode sites. These findings suggest that personalized beta stimulation effectively engages focal speech-motor network. In contrast, tRNS elicited broader and more robust modulatory effects by enhancing the engagement of task-relevant neural dynamics, specifically in the beta-band (15-30 Hz). The induced modulation in older adults was temporally and spectrally more relevant to the demanded task, implying that aging neural system is more susceptible to exogenous stimulation than that of younger adults. These findings can be interpreted from two perspectives. First, the frequency-specific stimulation (such as beta-tACS) entrains endogenous neural oscillations through phase alignment and enhances synchronization at the targeted frequency. Such stimulation is particularly suited to engaging stable oscillatory processes associated with specific functions (e.g., the alpha frequency during attention or beta oscillation during simple motor tasks). In contrast, in complex and multi-frequency tasks such as speech production, which rely on a more distributed network and rapid transitioning shifts across neural processing, the effects of frequency-specific stimulation may remain more spatially and spectrally focal than those induced by broadband stimulation such as tRNS.

Moreover, age-related decline in cognitive and motor functioning is associated with delayed neural communications [51, 52]. Compensatory increases in neural firing rate may partially mitigate these changes; however, they can cause variability in neural activity, leading to redistribution of spectral power and overlap of frequency bands [53]. Accordingly, a frequency-specific stimulation (i.e., beta-tACS) might be less efficient in entraining these diffused neural dynamics. In contrast to tACS, which is confined to a single frequency band, tRNS delivers a stochastic stimulation in which the frequency of the current varies in a random manner, enabling possible modulation of neural activity across broader frequency bands [54]. Hence, tRNS may be more beneficial to older adults due to its capacity to amplify subthreshold neural firing, increase cortical excitability, and engage distributed oscillatory patterns and large-scale cortical networks. This is particularly relevant in the context of speech production, a multi-layered process involving a larger network of cortical regions and several frequency bands [55, 56]. This finding is in line with the study of Inukai et al. (2016) comparing several tES techniques (tDCS, tACS, and tRNS) and reported that tRNS provided the most robust and consistent cortical excitability at multiple time points (immediately, at 5, 10, 15 and 20 minutes post-stimulation) [57]. Compared with tACS, stochastic resonance delivered through tRNS has shown the potential to enhance or disrupt neural synchrony depending on the underlying neural state; for instance, based on increased neural variability in aging or dysregulated oscillatory patterns in tinnitus patients [58].

In older adults, tRNS produced a more diffuse effect and modulated both alpha (9-13 Hz) and beta (15-30 Hz) bands across a broader set of electrode sites, including central, bilateral frontal, and right frontocentral sites for a more sustained period. This diffuse and multi-site modulation may also reflect recruitment of additional cortical areas to support task demands, compensating for reduced neural efficiency [59]. In contrast, younger adults exhibited a more localized and brief pre-stimulus modulation in beta-band (15-30 Hz), confined to the central electrodes. This pattern indicates the stabilization of motor activity and maintenance of current motor readiness state in the context of sequential response preparation, consistent with the involvement of the SMA in motor coordination and preparation [35, 60]. Furthermore, theta power (4-8 Hz) which is intricately associated with motor preparation and voluntary motor movement [61], showed a tRNS-induced decrease in the early preparatory phase across the frontal and central electrode sites in young adults. This modulation may account for a widespread reduction of demand for cognitive control and more automatic engagement in a simple and repetitive task, suggesting efficient neural processing.

Johari and Behroozmand (2020) reported a significant desynchronization of alpha and beta-band neural oscillations in older adults compared with younger adults during the planning phase of speech production [62]. Similar to their findings, our results demonstrated a stronger and more sustained alpha and beta power decrease in older participants reflecting increased neural efforts and compensatory mechanisms. Importantly, tRNS could potentially enhance such mechanisms and amplify the underlying oscillatory and compensatory mechanisms to facilitate the engagement of motor network during preparatory and execution phases. This pattern is also supported by the observed modulation induced by TtS, confined to right frontocentral electrodes, highlighting more selective and localized effects of personalized beta-tACS compared to the distributed effects of tRNS.

Although decreased alpha and beta-band power is likely associated with compensatory mechanisms, pronounced modulation induced by tRNS may reflect effective engagement of the motor network, supported by improved behavioral outcomes. Moreover, such modulation supports coordination of sensorimotor networks by reducing inhibitory control over task-related regions, which prepares the system to process relevant information. Consistent with previous literature, alpha desynchronization has been associated with enhanced neural excitability, functional inhibition of irrelevant information, and active processing [63]. Similarly, beta desynchronization reflects the active engagement of the motor network during preparation and execution [64]. Our findings are also in line with a prior study showing that alpha/beta desynchrony enhanced neural processing at the cortical level by reducing neuronal noise and improving signal-to-noise ratio [65].

## Conclusion

This study provides novel evidence that HD-tRNS and personalized HD-tACS were more effective than fixed-frequency HD-tACS in modulating the speech motor network at both behavioral and neural levels in older adults. Personalized tACS produced more focal effects, whereas tRNS induced broader modulation, engaging a distributed neural network by enhancing cortical excitability and recruiting different oscillatory frequencies.

These findings have implications for neurological disorders such as Parkinson’s disease or apraxia of speech with impaired motor control or altered neural dynamics. Broadband modulation of tRNS is a promising approach to restore an efficient neural network in speech production, working memory, complex motor tasks, and sensorimotor integration, which rely on multiple brain regions and oscillatory frequencies. Future studies are warranted to assess the long-term effects of the stimulation in clinical populations with impaired speech motor functioning in comparison with healthy older controls.

A limitation of this study was that our experimental paradigm was relatively simple and controlled. More complex speech tasks are required to capture multi-level (cognitive, linguistic, and motor) processes underlying naturalistic speech production.

## Data availability statement

The data that support the findings of this study are available from the authors upon reasonable requests.

## Source of funding

This study is financially supported by the Grant from Louisiana Board of Regent Research Competitiveness Program (Award Number: AWD-004500).

## Conflict of interests

The authors declare no conflict of interest.

## Ethical statement

This study was approved by the Institutional Review Board of the Louisiana State University (IRB approval Number: IRBAM-21-1019).

---

**Figure S1.**
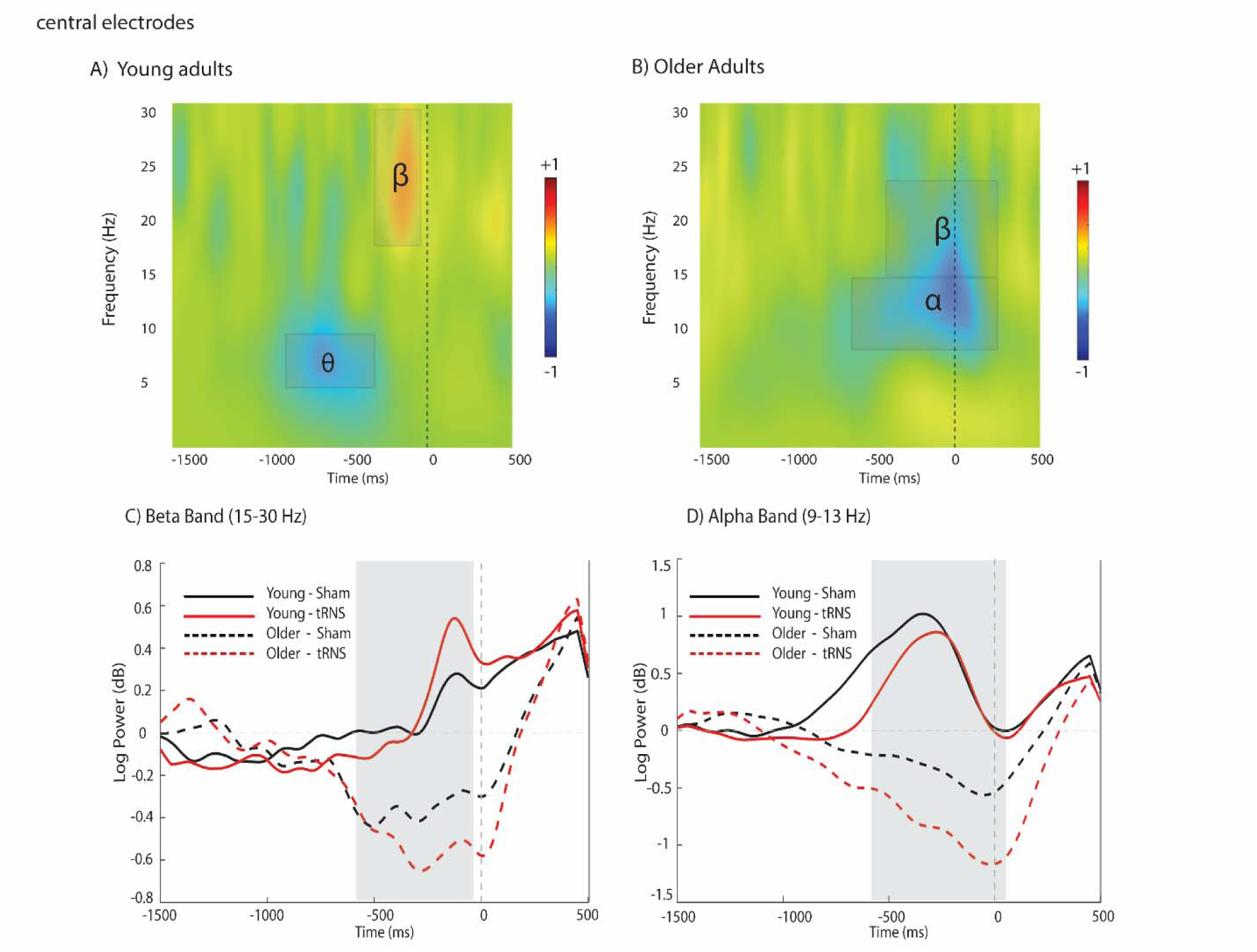
Neural activity following tRNS stimulation relative to Sham over the central electrodes (C1, Cz, C2) in young and older adults, across the time course of 1500 ms before to 500 ms after the onset of speech production. Time-frequency plots show power difference (tRNS—sham) for young (A) and older adults (B). The black rectangles highlight the alpha/beta-band time-frequency cluster of interest in older adults and theta/beta band time-frequency cluster of interest in young adults, and the vertical dashed line at 0 marks the onset of vocalizing the speech sound. Overlaid temporal profiles of log power (in decibels: dB) show the beta band (panel C) and the alpha band (panel D) time course for young and older adults in tRNS (red) and sham (black) conditions. Solid lines represent young adults, and dashed lines represent older adults. The shaded regions indicate the temporal clusters that reached significance in a cluster-based permutation analysis (between-group difference).

## Beta Band Power in tRNS-sham condition

Beta-band power was analyzed during speech production over the central electrodes (C1, Cz, C2) using cluster-based permutation analysis. No significant difference was observed between older and younger adults in the sham condition (p = 0.127); however, a significant group-difference cluster was identified spanning from −565 to -65 ms (p = 0.012) in the tRNS condition. Within-group comparisons showed no significant clusters between sham and tRNS in either the young or older adult groups group (p > 0.5).

Younger adults also showed a significant cluster of tRNS-sham difference in beta band around the stimulus onset (-660 to -565 ms) and in theta band (4-8 Hz) spanning -980 to -325 ms (p < 0.001). This modulation was not observed in older adults (p > 0.5).

## Alpha Band Power in tRNS-sham condition

The analysis of the alpha power over central electrodes (C1, Cz, C2) for the speech stimulus showed a significant difference in the sham condition between young and older participants from −765 to −90 ms (p = 0.014). Younger and older adults also differed significantly following tRNS stimulation within a cluster from −610 to 80 ms (p < 0.001). Within the older adult group, a significant cluster occurred between sham and tRNS from -180 to 100 ms (p < 0.001).

**Figure S2.**
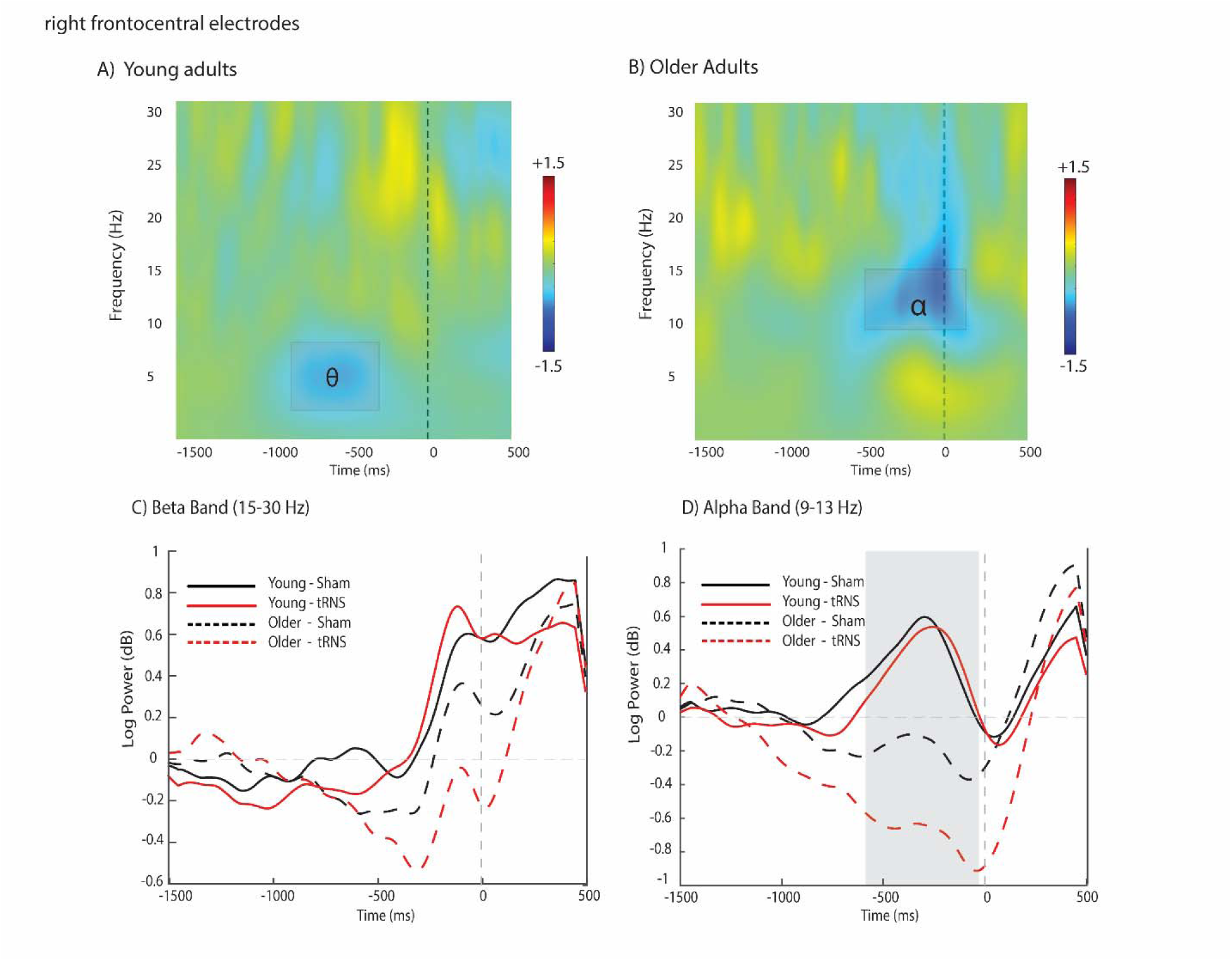
Neural activity following tRNS stimulation relative to Sham over right frontocentral electrodes (AF8, F8, F4, F6, FC6) in young and older adults, over the time course of 1500 ms before to 500 ms after the onset of speech production. Time-frequency plots show power difference (tRNS-sham) for young (A) and older adults (B). The black rectangles highlight the alpha-band time-frequency cluster of interest in older adults and the theta band (4-8 Hz) time-frequency cluster of interest in young adults, and the vertical dashed line at 0 marks the onset of vocalizing the speech sound. Overlaid temporal profiles of log power (in decibels: dB) show the beta band (panel C) and the alpha band (panel D) time course for young and older adults in tRNS (red) and sham (black) conditions. Solid lines represent young adults, and dashed lines represent older adults. The shaded regions indicate the temporal clusters that reached significance in a cluster-based permutation analysis (between-group difference).

## Beta Band Power in tRNS-sham condition

Cluster-based permutation analysis showed no significant differences in the beta band power (15-30 Hz) between young and older adult groups for tRNS (p = 0.0514) and sham (p = 0.5112) conditions over the right frontocentral electrodes (AF8, F8, F4, F6, FC6). In older adult group, a significant cluster was observed between -275 to -185 ms before speech production (p < 0.001) following tRNS compared to sham. In younger adults, a significant cluster was observed in tRNS-sham only in theta band (4-8 Hz) from -980 to -325 ms (p < 0.001).

## Alpha Band Power in tRNS-sham condition

The analysis of the alpha-band power within the right frontocentral ROI (AF8, F8, F4, F6, FC6) revealed no significant group differences for the sham condition. In contrast, a significant younger-older group difference was observed from -630 to -40 ms (p = 0.013) in tRNS condition. Within-group comparisons showed no significant sham-tRNS differences in the young group in the alpha band (9-13 Hz). However, the aging group exhibited a significant cluster spanning -565 to -375 ms (cluster p < 0.001).

## Notes

Conflict of Interest: None

### Competing Interest Statement

The authors have declared no competing interest.

## References

1. Lee, J. and H.-J. Kim, Normal Aging Induces Changes in the Brain and Neurodegeneration Progress: Review of the Structural, Biochemical, Metabolic, Cellular, and Molecular Changes. Frontiers in Aging Neuroscience, 2022. 14.

2. Chiappini, E., et al., You Are as Old as the Connectivity You Keep: Distinct Neurophysiological Mechanisms Underlying Age-Related Changes in Hand Dexterity and Strength. Archives of Medical Research, 2025. 56(1).

3. Ghasemian-Shirvan, E., et al., Age-related differences of motor cortex plasticity in adults: A transcranial direct current stimulation study. Brain Stimulation, 2020. 13(6): p. 1588–1599.

4. Dhamala, M., et al., Age-related slowing down in the motor initiation in elderly adults. Plos One, 2020. 15(9).

5. Myerson, J., S. Robertson, and S. Hale, Aging and Intraindividual Variability in Performance: Analyses of Response Time Distributions. Journal of the Experimental Analysis of Behavior, 2013. 88(3): p. 319–337.

6. Summerside, E.M., et al., Slowing of Movements in Healthy Aging as a Rational Economic Response to an Elevated Effort Landscape. J Neurosci, 2024. 44(15).

7. Mattay, V.S., et al., Neurophysiological correlates of age-related changes in human motor function. Neurology, 2002. 58(4): p. 630–635.

8. Scally, B., et al., Resting-state EEG power and connectivity are associated with alpha peak frequency slowing in healthy aging. Neurobiology of Aging, 2018. 71: p. 149–155.

9. Vysata, O., et al., Age-Related Changes in the Energy and Spectral Composition of EEG. Neurophysiology, 2012. 44(1): p. 63–67.

10. Saha, S., et al., Local homeostasis preserves global neural dynamics compensating for structural loss during human lifespan aging. Communications Biology, 2025. 8(1).

11. Hall, M.C., et al., Age-related alterations in alpha and beta oscillations support preservation of semantic processing in healthy aging. npj Aging, 2025. 11(1).

12. Park, J., et al., Age-related changes in neural oscillations vary as a function of brain region and frequency band. Front Aging Neurosci, 2025. 17: p. 1488811.

13. Darna, M., et al., Frontal theta oscillations and cognitive flexibility: Age-related modulations in EEG activity. Aging Brain, 2025. 8: p. 100142.

14. Rossiter, H.E., et al., Beta oscillations reflect changes in motor cortex inhibition in healthy ageing. NeuroImage, 2014. 91: p. 360–365.

15. Hübner, L., B. Godde, and C. Voelcker-Rehage, Older adults reveal enhanced task-related beta power decreases during a force modulation task. Behavioural Brain Research, 2018. 345: p. 104–113.

16. Zheng, X.Y. and V. Piai, Neural Oscillations in the Aging Brain Associated With Interference Control in Word Production. Neurobiology of Language, 2025. 6.

17. Summers, J.J., N. Kang, and J.H. Cauraugh, Does transcranial direct current stimulation enhance cognitive and motor functions in the ageing brain? A systematic review and meta-analysis. Ageing Research Reviews, 2016. 25: p. 42–54.

18. Lv, Y., et al., A meta-analysis of the effects of transcranial direct current stimulation combined with cognitive training on working memory in healthy older adults. Front Aging Neurosci, 2024. 16: p. 1454755.

19. Brown, G. and M. Brown, Transcranial electrical stimulation in neurological disease. Neural Regeneration Research, 2022. 17(10).

20. Matar, S.J., et al., Transcranial Direct-Current Stimulation May Improve Discourse Production in Healthy Older Adults. Frontiers in Neurology, 2020. 11.

21. Cattaneo, Z., A. Pisoni, and C. Papagno, Transcranial direct current stimulation over Broca’s region improves phonemic and semantic fluency in healthy individuals. Neuroscience, 2011. 183: p. 64–70.

22. Fertonani, A., et al., The timing of cognitive plasticity in physiological aging: a tDCS study of naming. Frontiers in Aging Neuroscience, 2014. 6.

23. Nomura, T. and H. Kirimoto, Anodal Transcranial Direct Current Stimulation Over the Supplementary Motor Area Improves Anticipatory Postural Adjustments in Older Adults. Front Hum Neurosci, 2018. 12: p. 317.

24. Krause, B. and R. Cohen Kadosh, Not all brains are created equal: the relevance of individual differences in responsiveness to transcranial electrical stimulation. Front Syst Neurosci, 2014. 8: p. 25.

25. Laakso, I., et al., Inter-subject Variability in Electric Fields of Motor Cortical tDCS. Brain Stimul, 2015. 8(5): p. 906–13.

26. Fabbrini, A., et al., Transcranial alternating current stimulation modulates cortical processing of somatosensory information in a frequency- and time-specific manner. NeuroImage, 2022. 254.

27. Lafleur, L.P., et al., No aftereffects of high current density 10 Hz and 20 Hz tACS on sensorimotor alpha and beta oscillations. Sci Rep, 2021. 11(1): p. 21416.

28. Tabari, F., et al., Personalized beta band HD-tACS over the left SMA improves speech and limb movement by modulating prefrontal delta oscillations in neurotypical young adults. Journal of Neural Engineering, 2025. 22(5).

29. Del Felice, A., et al., Personalized transcranial alternating current stimulation (tACS) and physical therapy to treat motor and cognitive symptoms in Parkinson’s disease: A randomized cross-over trial. Neuroimage Clin, 2019. 22: p. 101768.

30. Moret, B., et al., Transcranial random noise stimulation (tRNS): a wide range of frequencies is needed for increasing cortical excitability. Scientific Reports, 2019. 9(1).

31. Cui, J., et al., The effect of transcranial random noise stimulation (tRNS) over bilateral parietal cortex in visual cross-modal conflicts. Scientific Reports, 2025. 15(1).

32. van der Groen, O., et al., Using noise for the better: The effects of transcranial random noise stimulation on the brain and behavior. Neuroscience & Biobehavioral Reviews, 2022. 138.

33. Brambilla, M., et al., The Effect of Transcranial Random Noise Stimulation on Cognitive Training Outcome in Healthy Aging. Front Neurol, 2021. 12: p. 625359.

34. Tabari, F., et al., Effects of personalized vs. non-personalized neurostimulation protocols in improving speech and limb reaction times. Neuroscience, 2026. 596: p. 158–170.

35. Tabari, F. and K. Johari, Supplementary motor area: A promising neurostimulation target to improve speech production. J Commun Disord, 2026. 120: p. 106630.

36. del Campo-Vera, R.M., et al., Neuromodulation in Beta-Band Power Between Movement Execution and Inhibition in the Human Hippocampus. Neuromodulation: Technology at the Neural Interface, 2022. 25(2): p. 232–244.

37. Espenhahn, S., et al., Cortical beta oscillations are associated with motor performance following visuomotor learning. Neuroimage, 2019. 195: p. 340–353.

38. Brainard, D.H., The Psychophysics Toolbox. Spat Vis, 1997. 10(4): p. 433–6.

39. Delorme, A. and S. Makeig, EEGLAB: an open source toolbox for analysis of single-trial EEG dynamics including independent component analysis. Journal of Neuroscience Methods, 2004. 134(1): p. 9–21.

40. Pion-Tonachini, L., K. Kreutz-Delgado, and S. Makeig, The ICLabel dataset of electroencephalographic (EEG) independent component (IC) features. Data in Brief, 2019. 25.

41. Gramfort, A., MEG and EEG data analysis with MNE-Python. Frontiers in Neuroscience, 2013. 7.

42. Johari, K. and R. Behroozmand, Neural correlates of speech and limb motor timing deficits revealed by aberrant beta band desynchronization in Parkinson’s disease. Clinical Neurophysiology, 2021. 132(10): p. 2711–2721.

43. Delorme, A. and S. Makeig, EEGLAB: an open source toolbox for analysis of single-trial EEG dynamics including independent component analysis. J Neurosci Methods, 2004. 134(1): p. 9–21.

44. Ito, T., J.H. Coppola, and D.J. Ostry, Speech motor learning changes the neural response to both auditory and somatosensory signals. Scientific Reports, 2016. 6(1).

45. Maris, E. and R. Oostenveld, Nonparametric statistical testing of EEG- and MEG-data. Journal of Neuroscience Methods, 2007. 164(1): p. 177–190.

46. Johari, K., D.B. den Ouden, and R. Behroozmand, Effects of aging on temporal predictive mechanisms of speech and hand motor reaction time. Aging Clin Exp Res, 2018. 30(10): p. 1195–1202.

47. Frolov, N.S., et al., Age-related slowing down in the motor initiation in elderly adults. PLoS One, 2020. 15(9): p. e0233942.

48. Radecke, J.O., et al., Personalized alpha-tACS targeting left posterior parietal cortex modulates visuo-spatial attention and posterior evoked EEG activity. Brain Stimul, 2023. 16(4): p. 1047–1061.

49. Brevet-Aeby, C., et al., Three repeated sessions of transcranial random noise stimulation (tRNS) leads to long-term effects on reaction time in the Go/No Go task. Neurophysiol Clin, 2019. 49(1): p. 27–32.

50. Wach, C., et al., Effects of 10 Hz and 20 Hz transcranial alternating current stimulation (tACS) on motor functions and motor cortical excitability. Behav Brain Res, 2013. 241: p. 1–6.

51. van der Willik, K.D., et al., Trajectories of Cognitive and Motor Function Between Ages 45 and 90 Years: A Population-Based Study. J Gerontol A Biol Sci Med Sci, 2021. 76(2): p. 297–306.

52. Jones, A., et al., Limited Evidence to Review-Is There an Association Between Cognition and Upper Extremity Motor Reaction Time in Older Adults? NeuroSci, 2025. 6(3).

53. Hong, S.L. and G.V. Rebec, A new perspective on behavioral inconsistency and neural noise in aging: compensatory speeding of neural communication. Front Aging Neurosci, 2012. 4: p. 27.

54. Antal, A. and C.S. Herrmann, Transcranial Alternating Current and Random Noise Stimulation: Possible Mechanisms. Neural Plast, 2016. 2016: p. 3616807.

55. Whitford, T.J., et al., Gamma and Theta/Alpha-Band Oscillations in the Electroencephalogram Distinguish the Content of Inner Speech. eNeuro, 2025. 12(2).

56. Zioga, I., et al., Alpha and Beta Oscillations Differentially Support Word Production in a Rule-Switching Task. eneuro, 2024. 11(4).

57. Inukai, Y., et al., Comparison of Three Non-Invasive Transcranial Electrical Stimulation Methods for Increasing Cortical Excitability. Frontiers in Human Neuroscience, 2016. 10.

58. Vanneste, S., F. Fregni, and D. De Ridder, Head-to-Head Comparison of Transcranial Random Noise Stimulation, Transcranial AC Stimulation, and Transcranial DC Stimulation for Tinnitus. Front Psychiatry, 2013. 4: p. 158.

59. Taube, W. and B. Lauber, The ageing brain: Cortical overactivation - How does it evolve? J Physiol, 2026. 604(2): p. 813–828.

60. Bruijn, S.M., J.H. Van Dieën, and A. Daffertshofer, Beta activity in the premotor cortex is increased during stabilized as compared to normal walking. Frontiers in Human Neuroscience, 2015. 9.

61. Bland, B.H., et al., Amplitude, frequency, and phase analysis of hippocampal theta during sensorimotor processing in a jump avoidance task. Hippocampus, 2006. 16(8): p. 673–81.

62. Johari, K. and R. Behroozmand, Event-related desynchronization of alpha and beta band neural oscillations predicts speech and limb motor timing deficits in normal aging. Behav Brain Res, 2020. 393: p. 112763.

63. Başar, E. and B. Güntekin, A short review of alpha activity in cognitive processes and in cognitive impairment. International Journal of Psychophysiology, 2012. 86(1): p. 25–38.

64. Pierrieau, E., et al., Changes in cortical beta power predict motor control flexibility, not vigor. Communications Biology, 2025. 8(1).

65. Griffiths, B.J., et al., Alpha/beta power decreases track the fidelity of stimulus-specific information. Elife, 2019. 8.

